## Supplementary File 1 for "MicroRNA-like snoRNA-derived RNAs (sdRNAs) promote castration resistant prostate cancer"

**Supplementary File 1. Select misannotated human microRNAs processed from snoRNAs.**

Images generated by Ensembl genome browser. **(A)** Human microRNA hsa-miR-664b is entirely embedded in SNORA36A

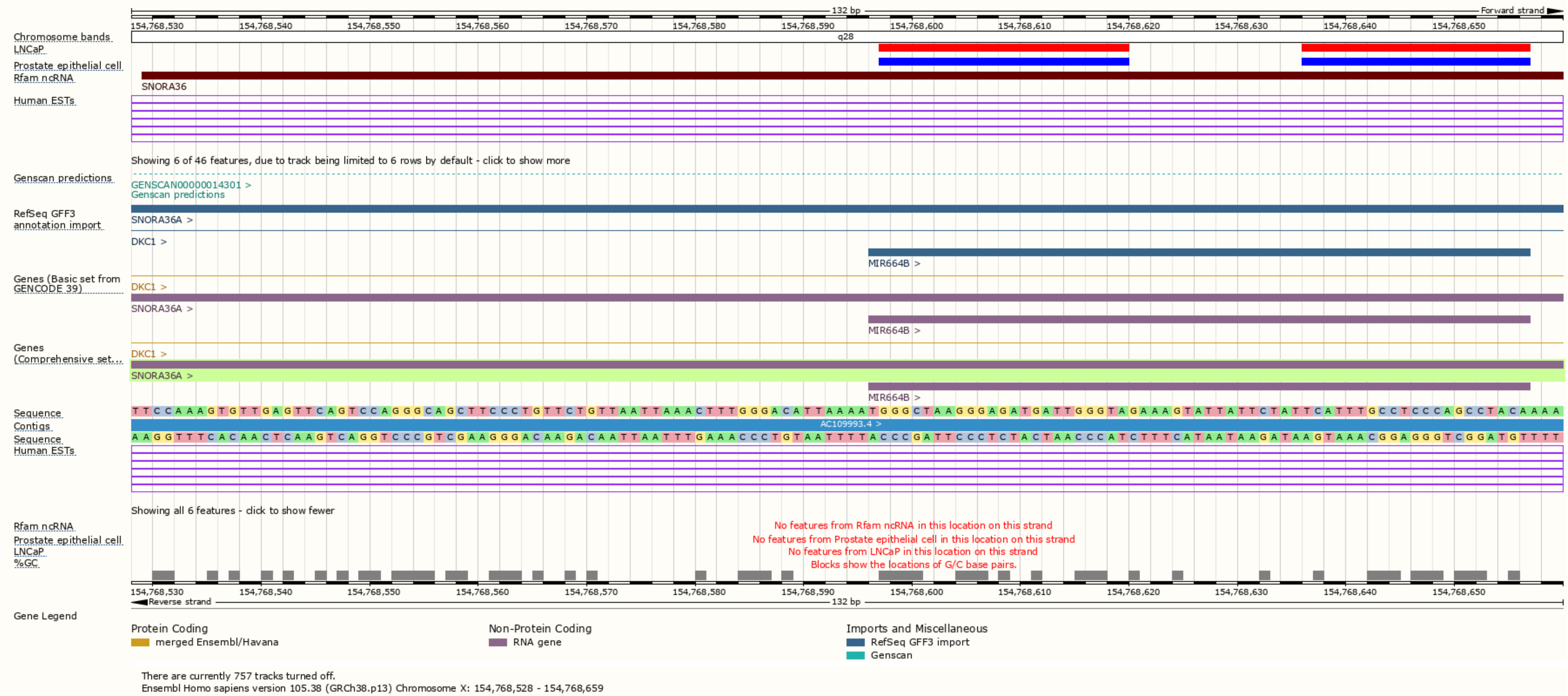

**Supplementary File 1. Select misannotated human microRNAs processed from snoRNAs.**

**(B)** Human microRNA hsa-miR-664a is entirely embedded in SNORA36B.

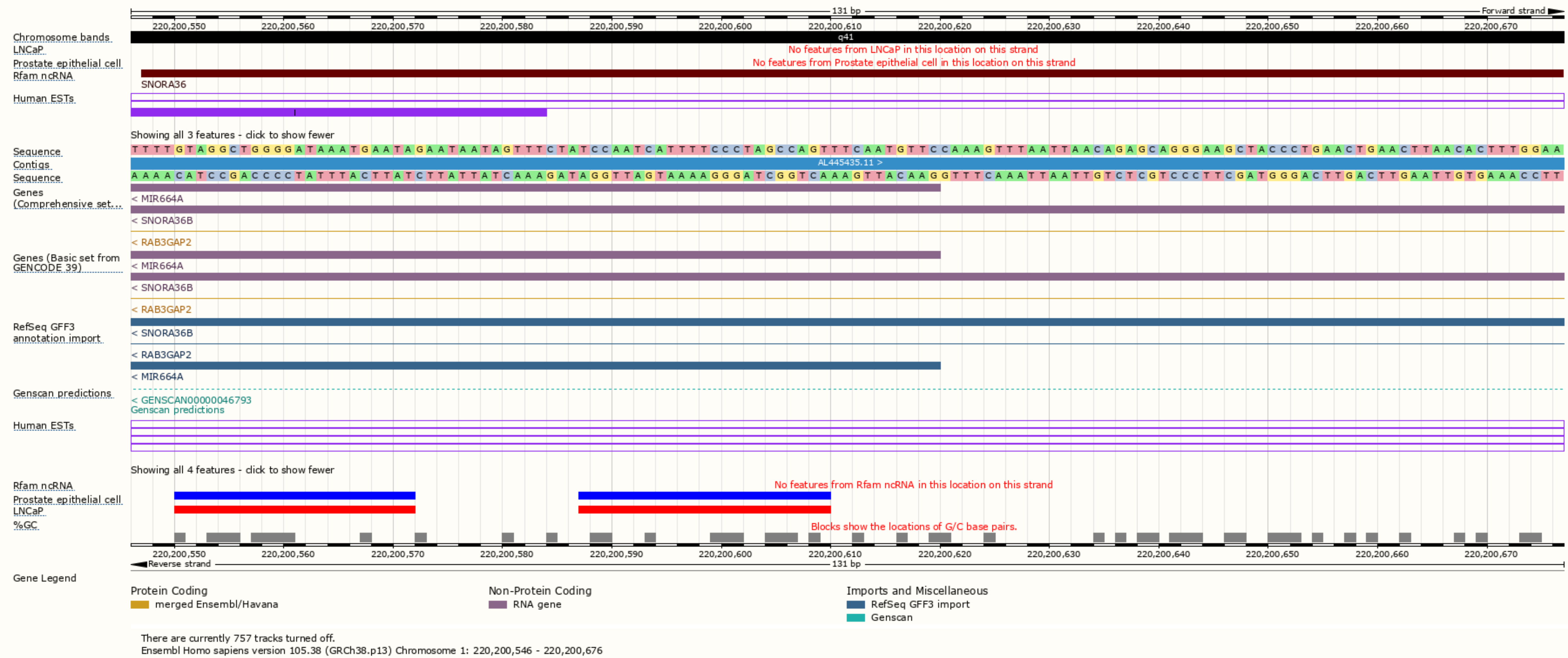

**Supplementary File 1. Select misannotated human microRNAs processed from snoRNAs.**

**(C)** Human microRNA hsa-miR-1291 is entirely embedded in SNORA2C.

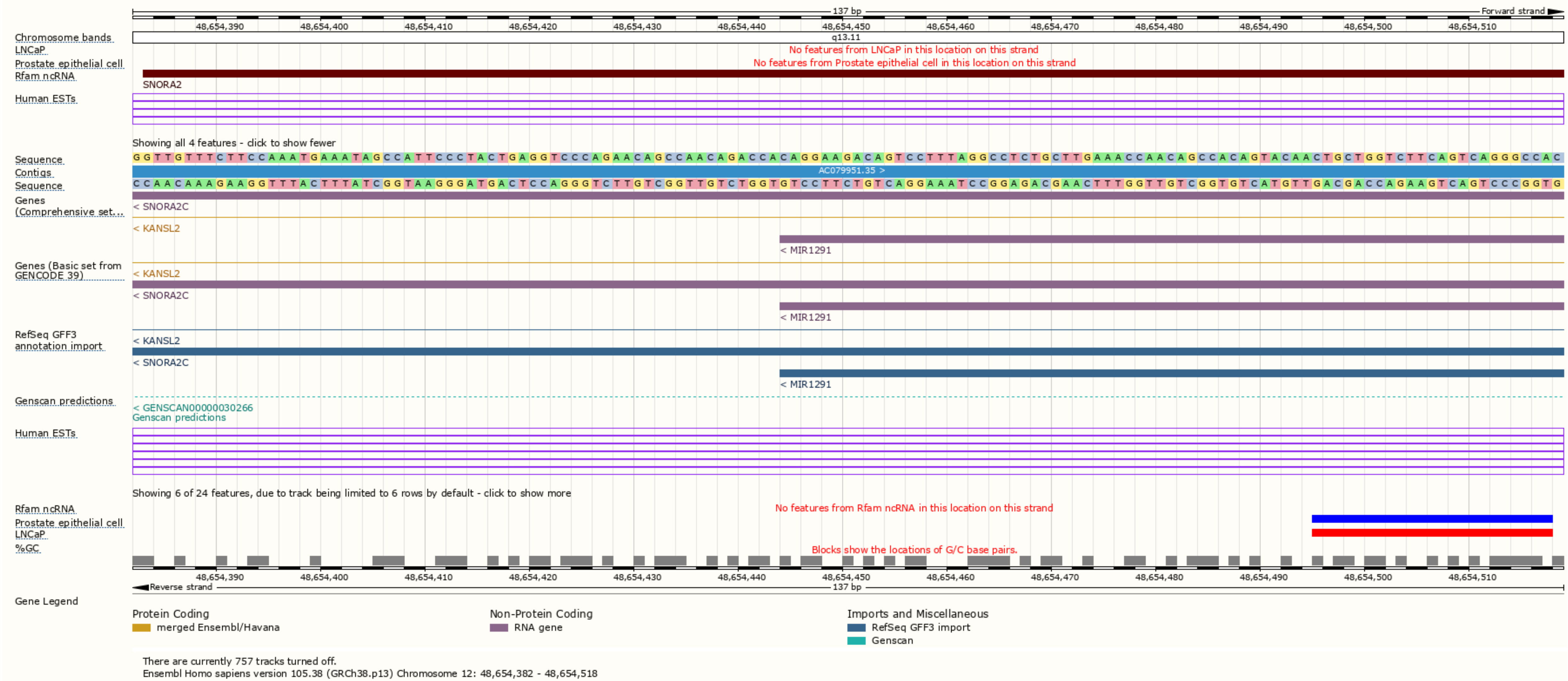

**Supplementary File 1. Select misannotated human microRNAs processed from snoRNAs.**

**(D)** Human microRNA hsa-miR-3651 is entirely embedded in SNORA84.

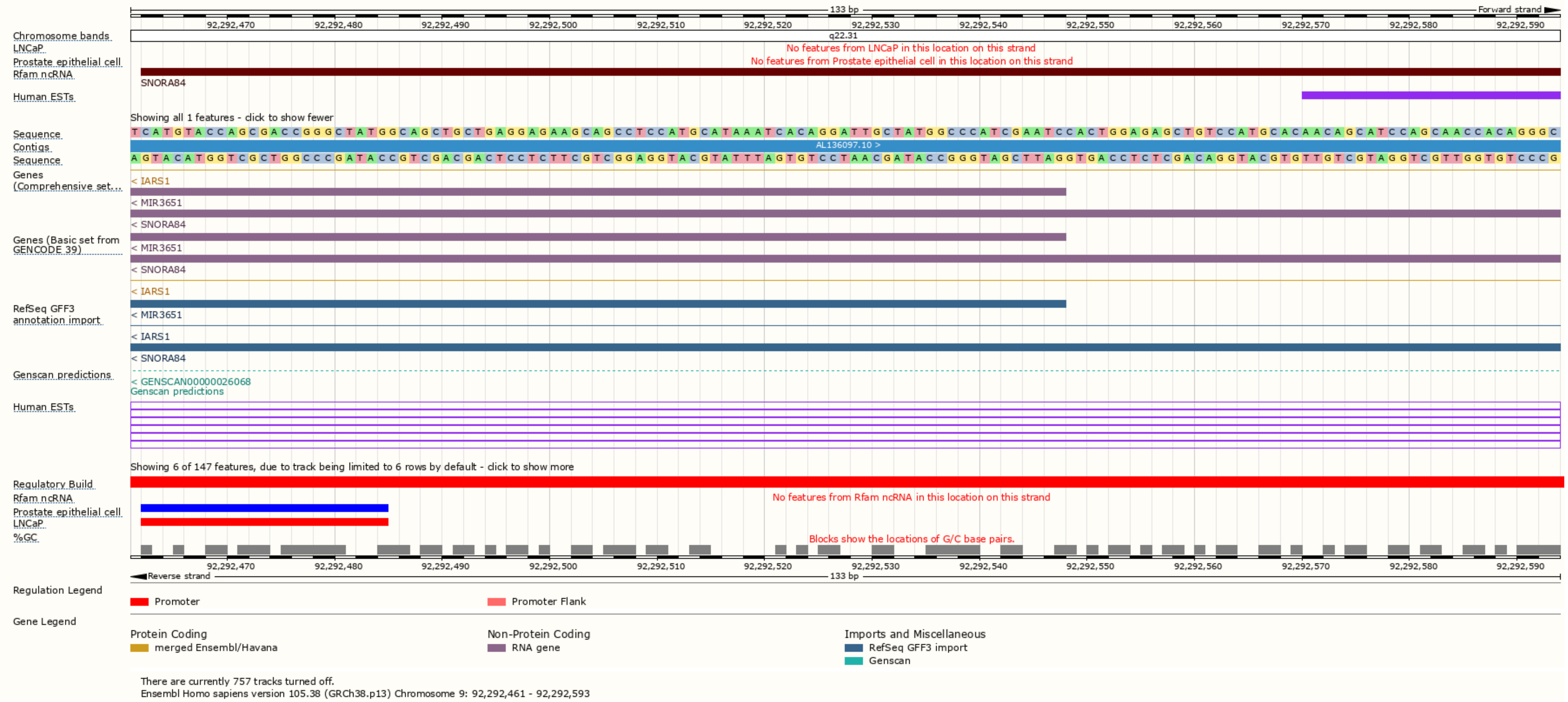

**Supplementary File 1. Select misannotated human microRNAs processed from snoRNAs.**

**(E)** Human microRNA hsa-miR-1248 is entirely embedded in SNORA81.

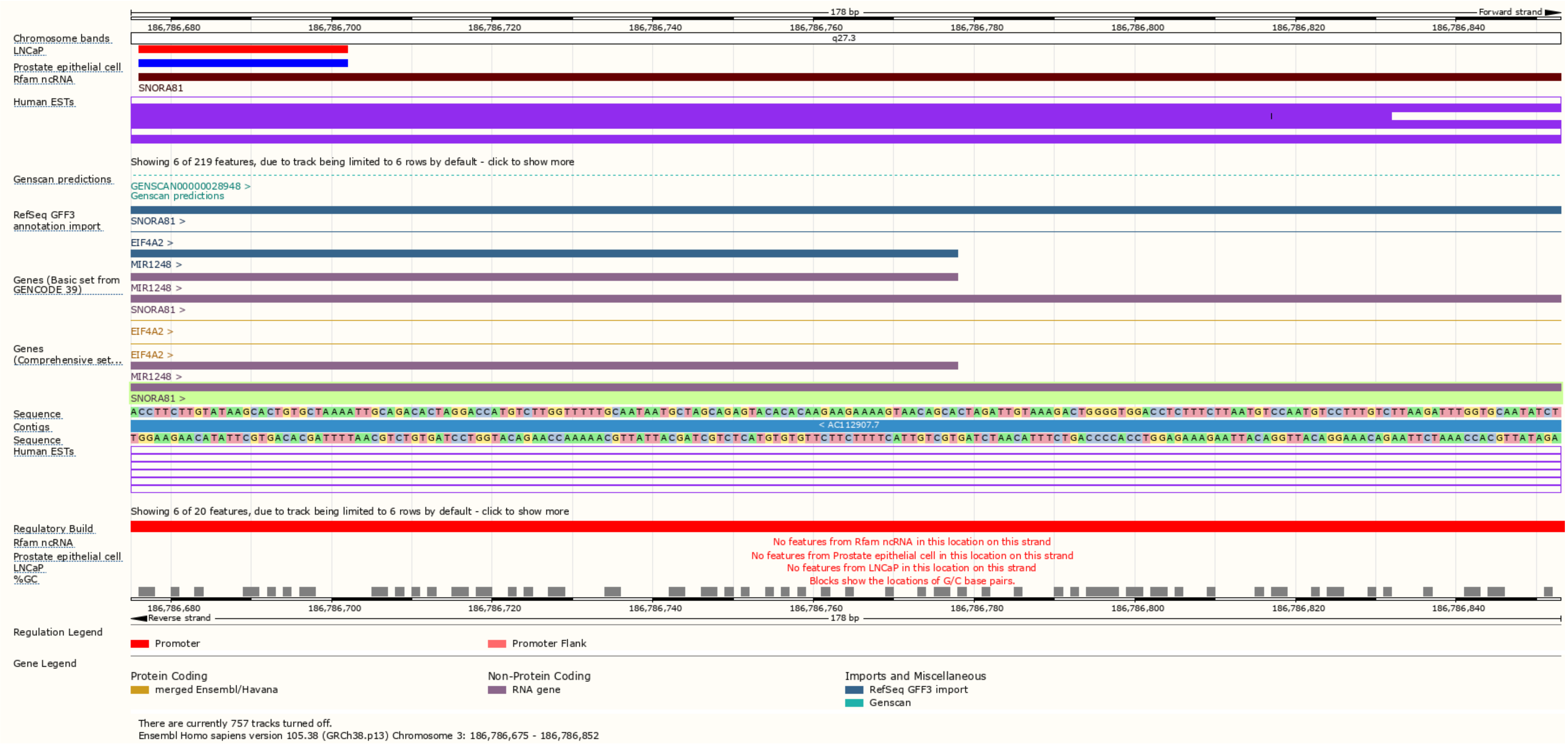
