## Supplementary File 2 for "MicroRNA-like snoRNA-derived RNAs (sdRNAs) promote castration resistant prostate cancer"

**Supplementary File 2. 38 SURFR-identified sdRNAs significantly differentially expressed in TCGA PRAD tumor samples as compared to control prostate tissues**All samples were acquired from The Cancer Genome Atlas (TCGA) Research network PRAD dataset and are publicly available at https://www.cancer.gov/tcga. SURFR analysis of TCGA PRAD and normal prostate control returned expression in reads per million (RPM) for each sdRNA detected. Rstudio was used to calculate differential expression and rank each sdRNA by cancer prevalence (% of TCGA samples that expressed the sdRNA) and differential expression. Significant results were constricted to those sdRNAs with ≥ 2x fold change in prostate cancer and were expressed at ≥ 30 RPM in a minimum of 50% of TCGA PRAD small RNA-seq files. "< 30*" indicates that a RPM was determined to be less than 30 but set to 30 for the purpose of differential expression calculation due to the inherent 30 RPM cutoff during SURFR analysis.

| Ensembl Gene ID | SnoRNA | Start Position | Stop Position | Length | Cancer Average Reads per Million (RPM) | Cancer File Count (/489) | Control Average Reads per Million (RPM) | Control File Count (/52) | Cancer/Control Fold Change | Sequence (5' to 3') |
|---|---|---|---|---|---|---|---|---|---|---|
| ENSG00000206754 | SNORD101 | 3 | 26 | 24 | 185 | 444 | < 30* | 0 | 6.17 | UUGAAUGAUGACUUUAAUUGUCGG |
| ENSG00000275043 | SNORD25 | 43 | 65 | 23 | 1312 | 470 | 235 | 42 | 5.59 | CGUGAGGAUAAAUAACUCUGAGG |
| ENSG00000199753 | SNORD104 | 45 | 70 | 26 | 10102 | 489 | 1896 | 52 | 5.33 | CGGGUGAUGCGAACUGGAGUCUGAGC |
| ENSG00000201823 | SNORD48 | 38 | 61 | 24 | 774 | 282 | 150 | 14 | 5.18 | UGAUGCCAUCACCGCAGCGCUCUG |
| ENSG00000264549 | SNORD95 | 39 | 62 | 24 | 1623 | 448 | 333 | 52 | 4.88 | UGCUGAAAUCCAGAGGCUGUUUCU |
| ENSG00000275994 | SNORA24 | 1 | 25 | 25 | 711 | 477 | 150 | 16 | 4.75 | CUCCAUGUAUCUUUGGGACCUGUCA |
| ENSG00000207280 | SNORD20 | 2 | 24 | 23 | 628 | 486 | 136 | 26 | 4.61 | GGAUAUGAUGACUGAUUACCUGA |
| ENSG00000283551 | SNORD98 | 40 | 64 | 25 | 1874 | 488 | 428 | 52 | 4.37 | GCAGUGUGGAACACAAUGAACUGAA |
| ENSG00000275996 | SNORD27 | 2 | 22 | 21 | 598 | 267 | 144 | 23 | 4.16 | CUCCAUGAUGAACACAAAAUG |
| ENSG00000275996 | SNORD27 | 47 | 68 | 22 | 1233 | 483 | 305 | 52 | 4.05 | GUGAUGUCAUCUUACUACUGAG |
| ENSG00000276314 | SNORD107 | 2 | 29 | 28 | 129 | 255 | 34 | 3 | 3.79 | GUUCAUGAUGACACAGGACCUUGUCUGA |
| ENSG00000199477 | SNORA31 | 2 | 24 | 23 | 1470 | 393 | 395 | 16 | 3.72 | UGCAUCCACUGAUAGACCUUGAA |
| ENSG00000212452 | SNORD69 | 26 | 48 | 23 | 479 | 362 | 129 | 10 | 3.72 | GGAUCUGACUGACUGUGCUGAGU |
| ENSG00000226572 | SNORD57 | 2 | 25 | 24 | 420 | 469 | 113 | 23 | 3.71 | GGAGGUGAUGAACUGUCUGAGCCU |
| ENSG00000206630 | SNORD60 | 2 | 26 | 25 | 1350 | 489 | 376 | 49 | 3.59 | GUCUGUGAUGAAUUGCUUUGACUUC |
| ENSG00000206622 | SNORA69 | 112 | 131 | 20 | 336 | 346 | 98 | 5 | 3.42 | GACAGAUUGACAUGGACAAU |
| ENSG00000281859 | SNORD38B | 46 | 65 | 20 | 102 | 414 | < 30* | 0 | 3.40 | GAAGAUAAAGUGUGUCUGAG |
| ENSG00000221539 | SNORD99 | 49 | 68 | 20 | 415 | 371 | 131 | 3 | 3.16 | AUGCGGAUAUGGGACUGAGA |
| ENSG00000238917 | SNORD10 | 121 | 140 | 20 | 364 | 416 | 119 | 13 | 3.06 | UCAGUCUUUGUACUCUGAGA |
| ENSG00000221116 | SNORD110 | 53 | 73 | 21 | 299 | 449 | 100 | 15 | 3.00 | GAUGUCUCCAUGUCUCUGAGC |
| ENSG00000235408 | SNORA71B | 141 | 160 | 20 | 277 | 280 | 93 | 5 | 2.99 | UCUGGAGCUUUCGUACAUGC |
| ENSG00000263934 | SNORD3A | 196 | 215 | 20 | 3494 | 484 | 1205 | 43 | 2.90 | GAGAGAACGCGGUCUGAGUG |
| ENSG00000206979 | SNORD61 | 52 | 71 | 20 | 215 | 260 | 77 | 5 | 2.80 | UCCUCUAAGAAGUUCUGAGC |
| ENSG00000202031 | SNORD38A | 1 | 19 | 19 | 499 | 489 | 185 | 46 | 2.70 | UCUCGUGAUGAAAACUCUG |
| ENSG00000239039 | SNORD13 | 83 | 104 | 22 | 252 | 325 | 94 | 7 | 2.67 | UGGGCACAUUACCCGUCUGACC |
| ENSG00000206989 | SNORD63 | 40 | 61 | 22 | 293 | 480 | 110 | 45 | 2.67 | AACGUGUGGAAAACUAAUGACU |
| ENSG00000239043 | SNORD127 | 5 | 27 | 23 | 217 | 264 | 84 | 2 | 2.58 | AACUGUGAUGAAAGAUUUGGUCU |
| ENSG00000266300 | SNORD52 | 38 | 64 | 27 | 276 | 362 | 109 | 4 | 2.53 | GGUCAUGAUGUCAAAACUAAGUUCUGA |
| ENSG00000263764 | SNORD43 | 2 | 22 | 21 | 395 | 484 | 157 | 40 | 2.52 | ACAGAUGAUGAACUUAUUGAC |
| ENSG00000238942 | SNORD2 | 31 | 52 | 22 | 285 | 374 | 115 | 48 | 2.48 | CUGACCUGAAAUGAAGAGAAUA |
| ENSG00000264591 | SNORD84 | 39 | 66 | 28 | 638 | 417 | 260 | 35 | 2.45 | CGCAGUGAUGACCCUCAUCUAUCACCCU |
| ENSG00000249020 | SNORA58 | 114 | 134 | 21 | 254 | 378 | 108 | 11 | 2.37 | AUUGCAGGACUCUAAACAUUU |
| ENSG00000238862 | SNORD19B | 55 | 73 | 19 | 384 | 448 | 163 | 22 | 2.36 | UACAAGAUCCAACUCUGAU |
| ENSG00000238649 | SNORD42A | 41 | 60 | 20 | 281 | 425 | 124 | 13 | 2.27 | UGAACAAAGGAACCACUGAA |
| ENSG00000207031 | SNORD59A | 50 | 71 | 22 | 164 | 414 | 73 | 24 | 2.25 | GAAGCCACAUUUAGGUACUGAG |
| ENSG00000206656 | SNORD116-17 | 72 | 91 | 20 | 639 | 324 | 286 | 6 | 2.24 | AUCCUCGUCGAACUGAGGUC |
| ENSG00000206680 | SNORD21 | 71 | 90 | 20 | 374 | 466 | 175 | 44 | 2.14 | GUUUCAAGACGGGACUGAUG |
| ENSG00000212447 | SNORD90 | 79 | 97 | 19 | 872 | 390 | 416 | 25 | 2.10 | CCUACUGUGGAAUCUGAAG |
