## Supplementary File 3 for "MicroRNA-like snoRNA-derived RNAs (sdRNAs) promote castration resistant prostate cancer"

**Supplementary File 3. SdRNA -D19b and -A24 PC3 cell scratch assays.** Representative migration (wound-healing) assays for PC3 cells transfected with the indicated sdRNA mimic or inhibitor. **(A)** 19 inhibitor.

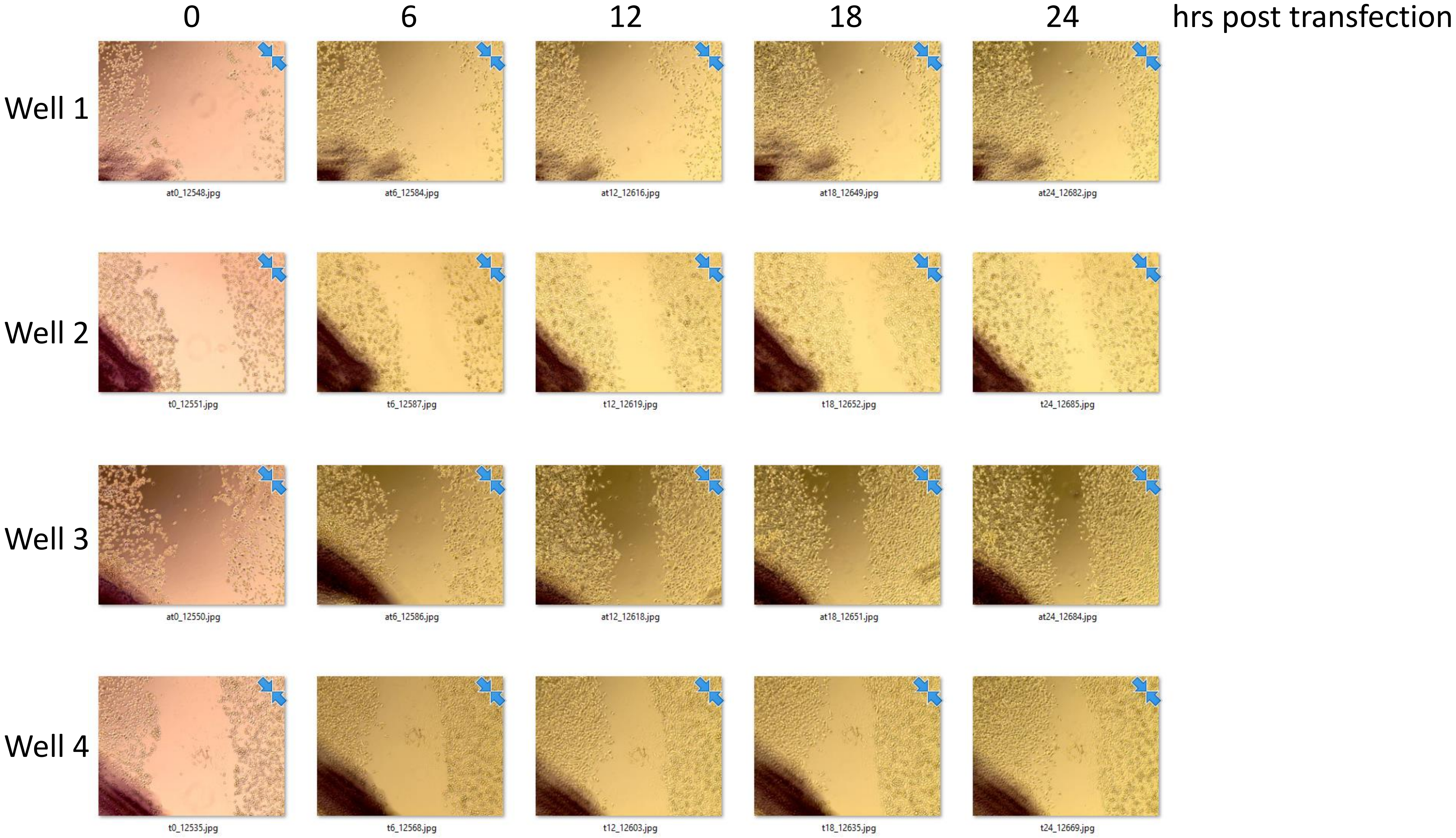

**Supplementary File 3. SdRNA -D19b and -A24 PC3 cell scratch assays. (B)** 19 mimic.

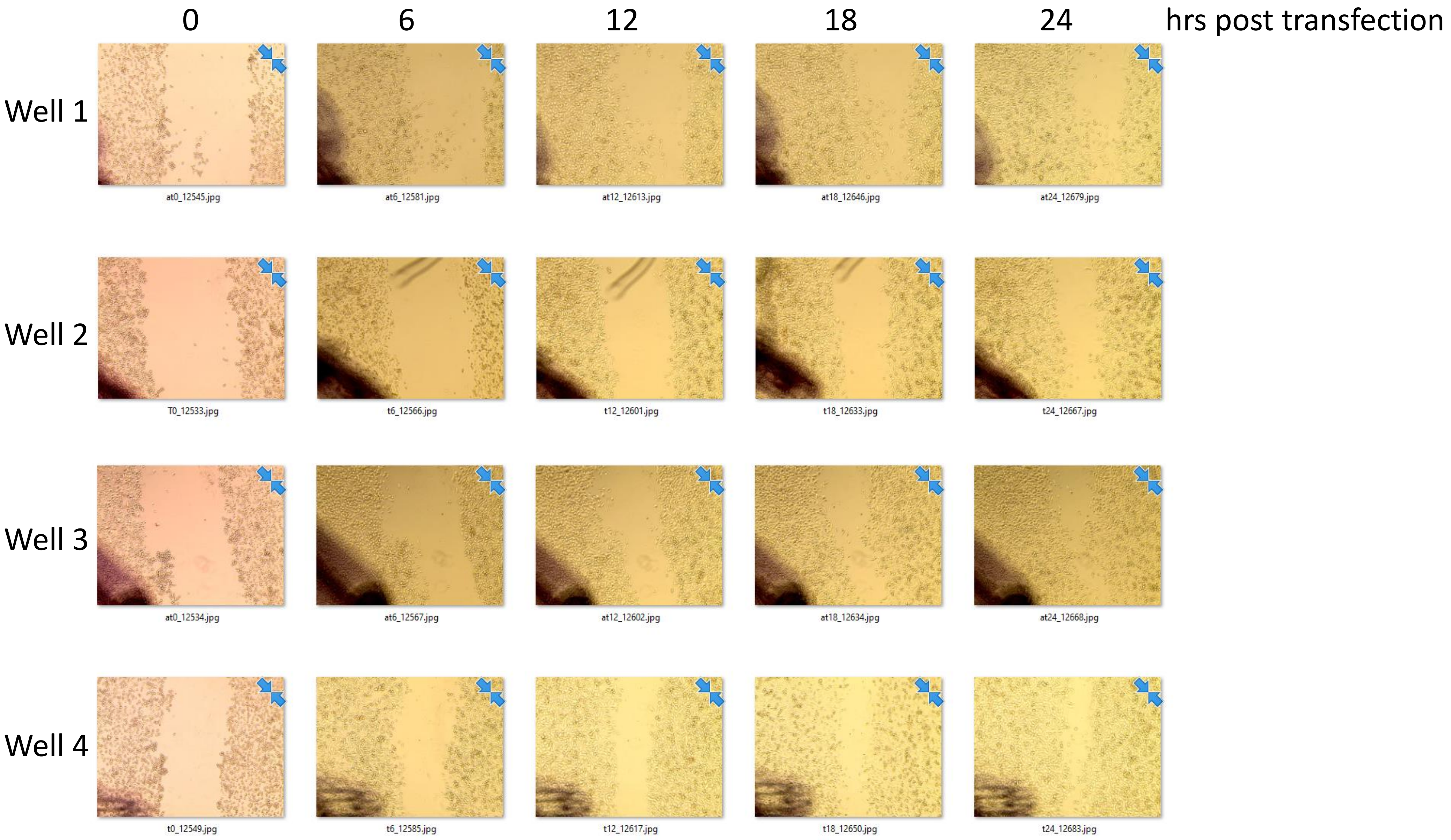

**Supplementary File 3. SdRNA -D19b and -A24 PC3 cell scratch assays. (C)** 24 inhibitor.

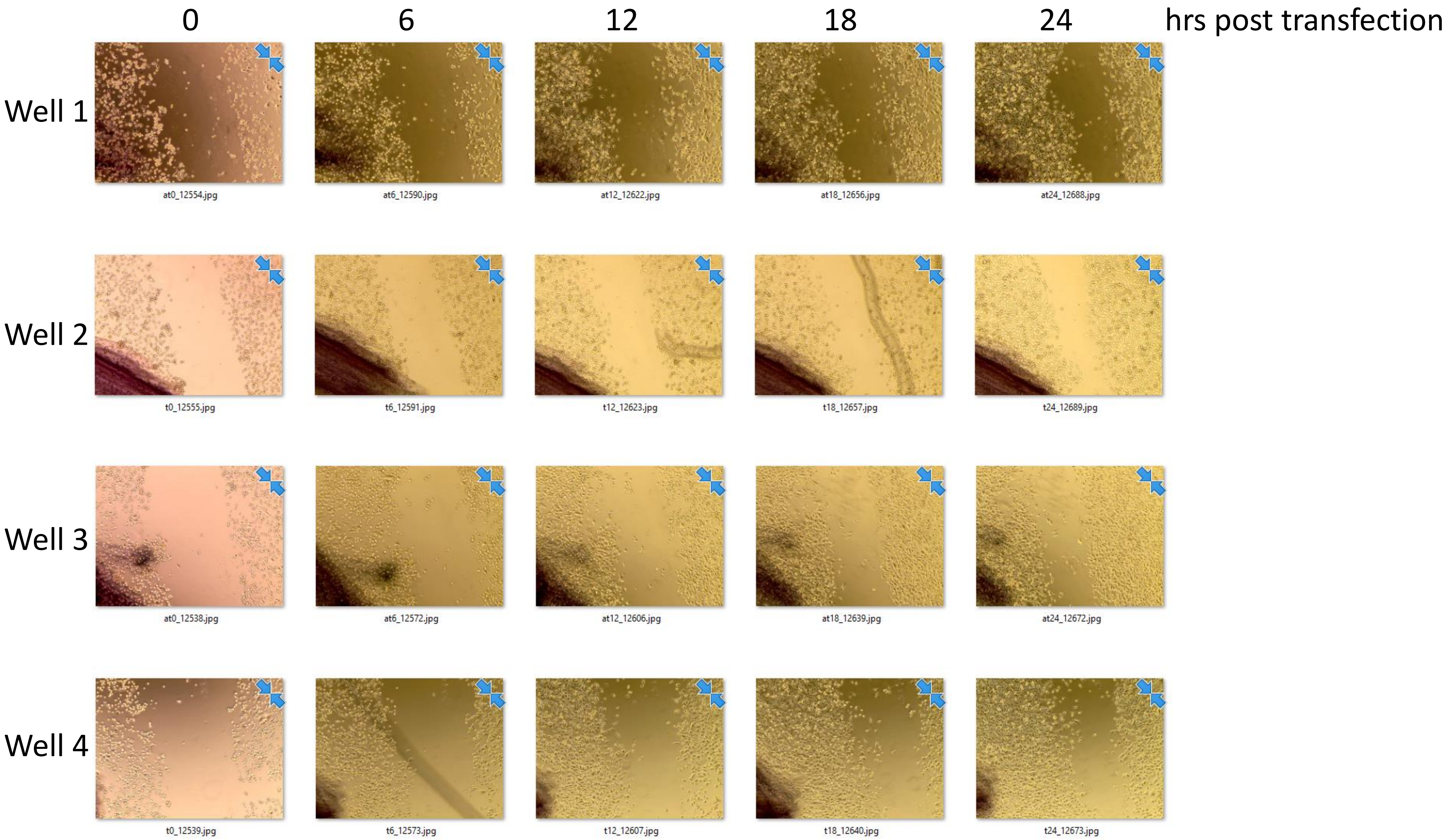

**Supplementary File 3. SdRNA -D19b and -A24 PC3 cell scratch assays. (D)** 24 mimic.

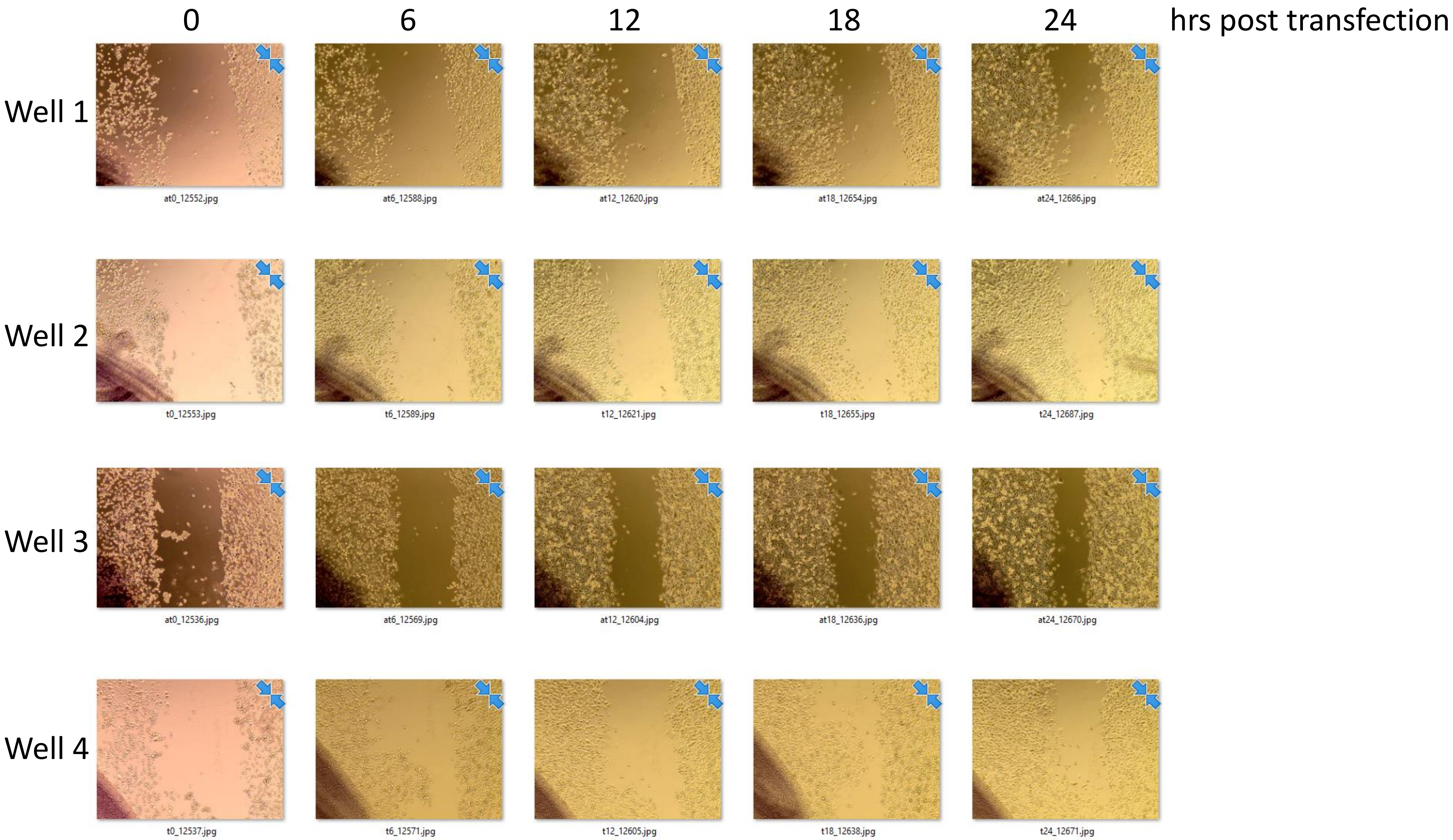

**Supplementary File 3. SdRNA -D19b and -A24 PC3 cell scratch assays. (E)** 42 inhibitor.

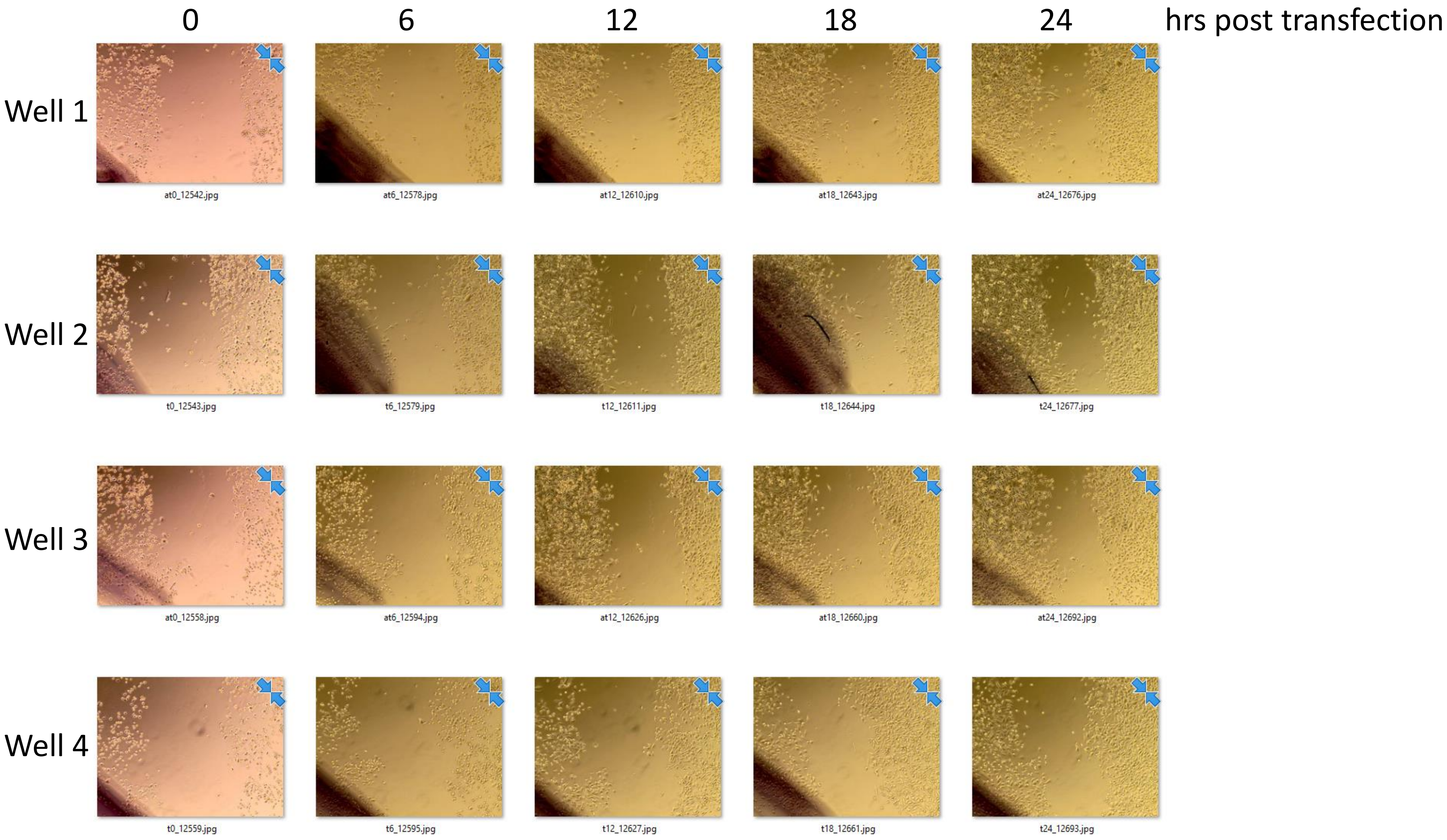

**Supplementary File 3. SdRNA -D19b and -A24 PC3 cell scratch assays. (F)** 42 mimic.

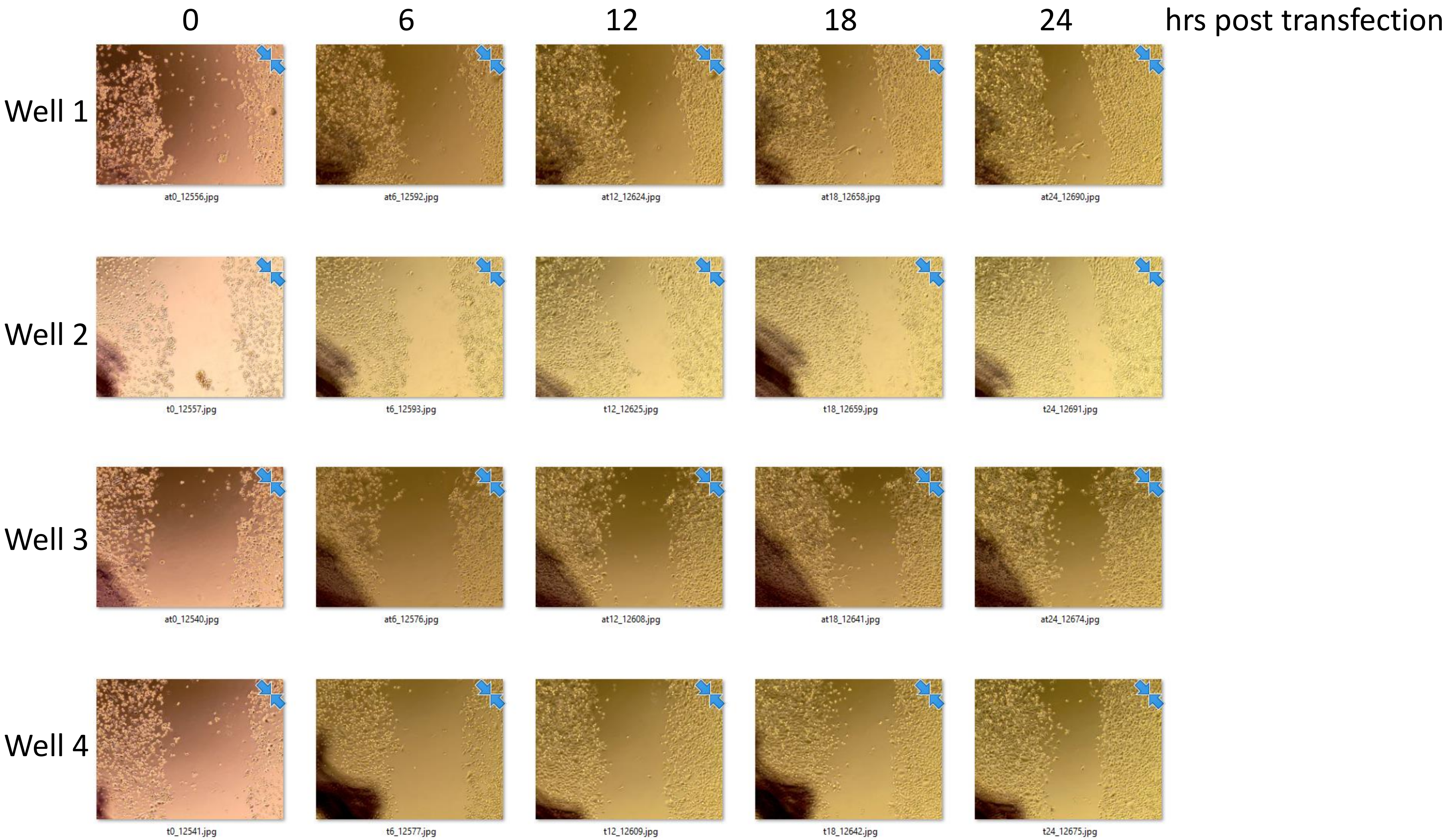

**Supplementary File 3. SdRNA -D19b and -A24 PC3 cell scratch assays. (G)** Control inhibitor.

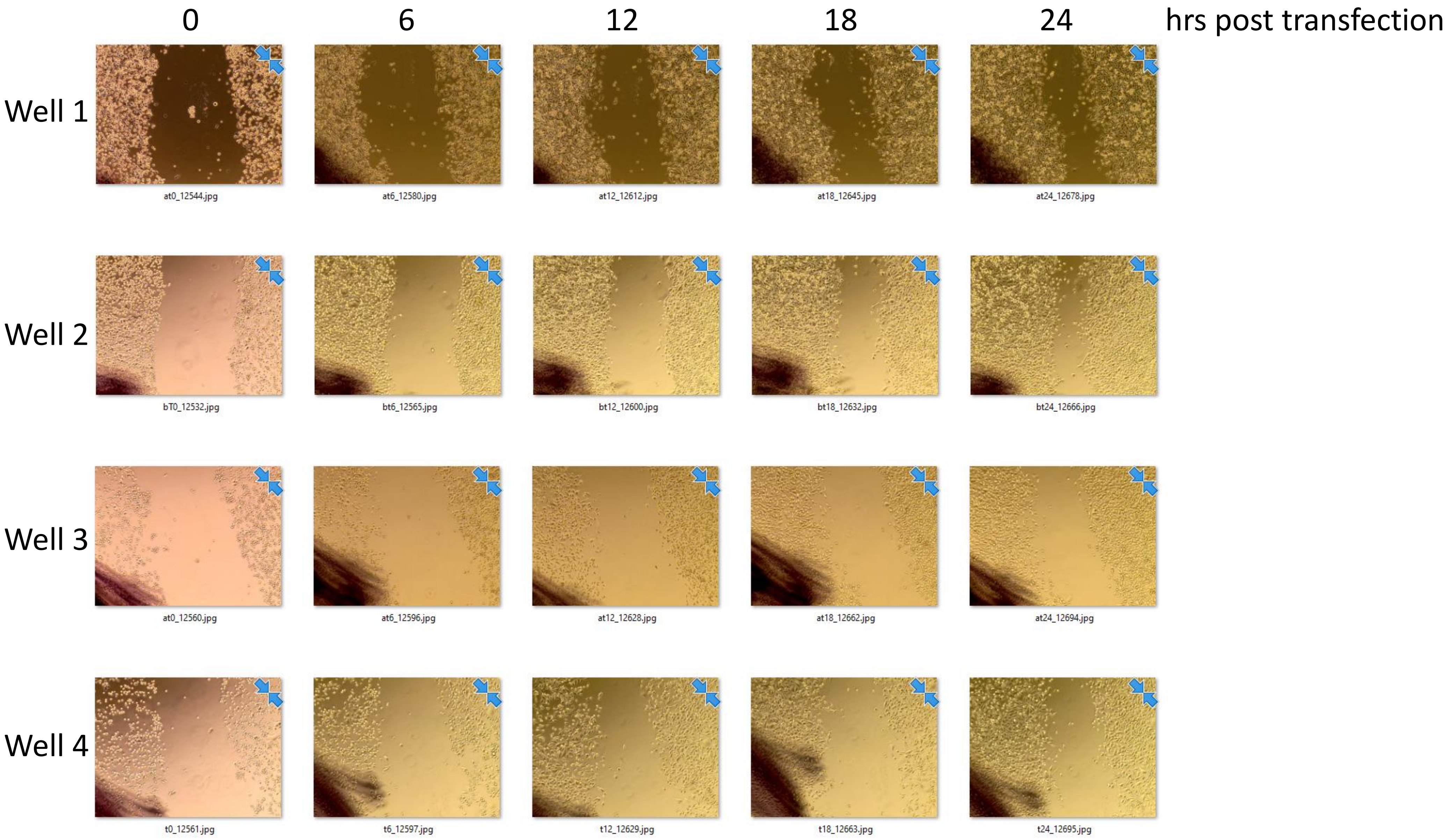

**Supplementary File 3. SdRNA -D19b and -A24 PC3 cell scratch assays. (H)** Control mimic.

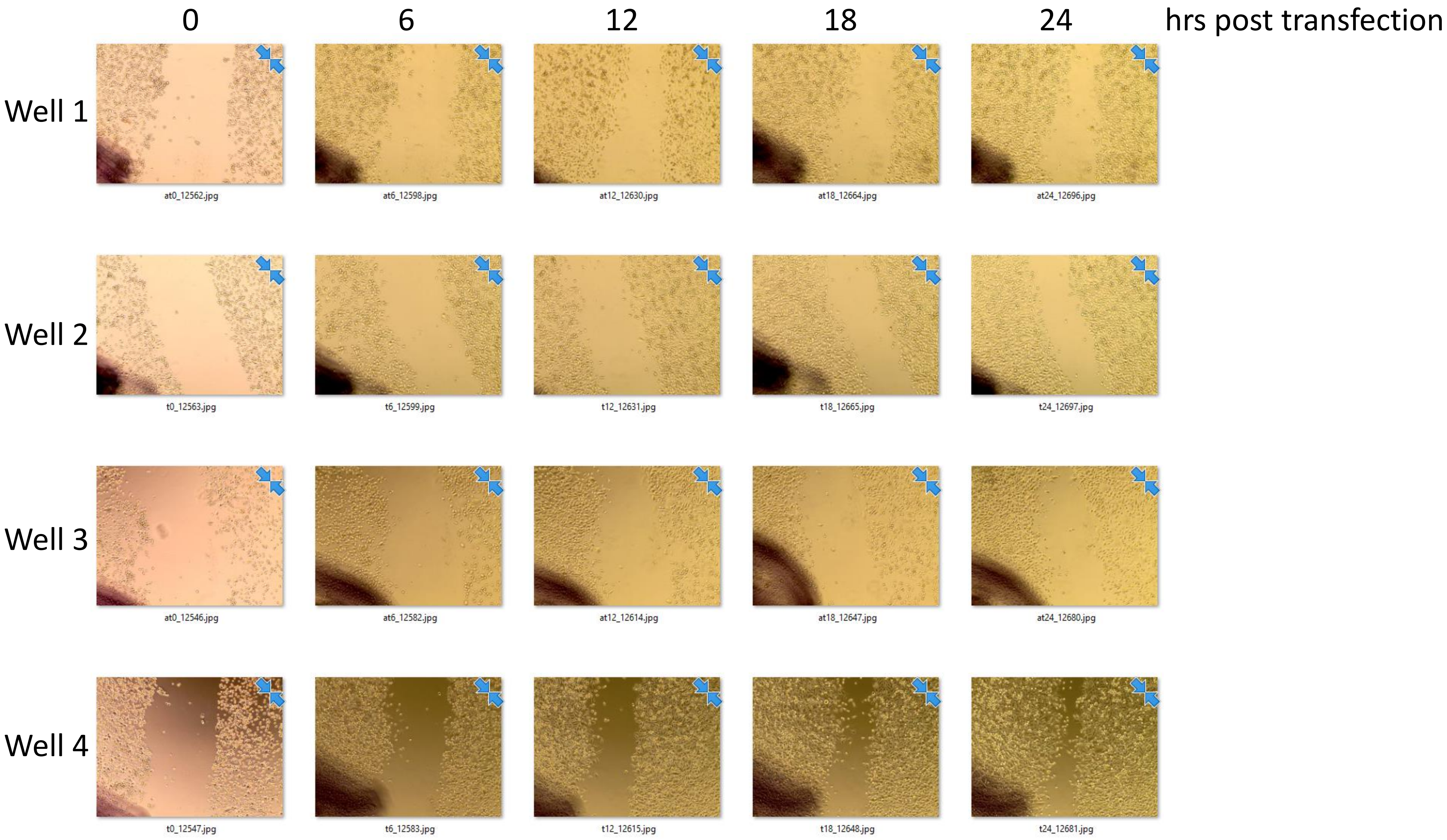
