## Supplementary File 5 for "MicroRNA-like snoRNA-derived RNAs (sdRNAs) promote castration resistant prostate cancer"

**Supplementary File 5. Oligonucleotide master list.** All sequences listed 5' to 3'.

**Mimics** were ordered as custom miRIDIAN miRNA mimics (Horizon Discovery) at 0.015 µmol scale. miRIDIAN Mimics are double-stranded RNA oligonucleotides chemically enhanced with the ON-TARGET modification pattern to preferentially program RISC with the active microRNA strand.

| | | |
|---|---|---|
| sd_A24 | Active strand | P-CUCCAUGUAUCUUUGGGACCUGUCA |
| | Passenger | ACAGGUCCCAAAGAUACAUGGAGUU |
| sd_D19b | Active strand | P-AUUACAAGAUCCAACUCUGAU |
| | Passenger | CAGAGUUGGAUCUUGUAAUUU |
| sd_D58c | Active strand | P-UUGCUGUGAUGACUAUCUUAGGAC |
| | Passenger | CCUAAGAUAGUCAUCACAGCAAUU |
| sd_D42A | Active strand | P-CAUGAACAAAGGAACCACUGAA |
| | Passenger | CAGUGGUUCCUUUGUUCAUGUU |
| sd_D24 | Active strand | P-CUAUCUGAGAGAUGGUGAUGAC |
| | Passenger | CAUCACCAUCUCUCAGAUAGUU |
| sd_Ctl | Active strand | P-UAAUCUGACCAAGACUCUUAA |
| | Passenger | AAGAGUCUUGGUCAGAUUAUU |

**Inhibitors** were ordered as custom IDT miRNA Inhibitors (Integrated DNA Technologies) at 5 nmol scale. IDT miRNA Inhibitors are RNA oligonucleotides comprised of 2'-O-methyl residues that confer increased binding affinity to RNA targets and resistance to endonuclease degradation. ZEN modifications are included to block exonuclease degradation.

| | |
|---|---|
| sdRNA-D19b | mA/ZEN/mUmCmAmGmAmGmUmUmGmGmAmUmCmUmUmGmUmAmA/3ZEN/ |
| sdRNA-A24 | mA/ZEN/mCmAmGmGmUmCmCmCmAmAmAmGmAmUmAmCmAmUmGmGmAmG/3ZEN/ |
| sdRNA-A42 | mC/ZEN/mAmGmUmGmGmUmUmCmCmUmUmUmGmUmUmCmAmUmG/3ZEN/ |
| sdRNA-A61 | mC/ZEN/mCmUmGmUmCmUmGmAmAmAmCmUmAmGmCmCmCmAmCmAmU/3ZEN/ |
| sdRNA-93 | mA/ZEN/mGmAmGmUmUmCmUmCmAmUmCmCmUmUmGmGmCmCmA/3ZEN/ |
| Scram Ctl | mA/ZEN/mAmGmAmGmUmCmUmUmGmGmUmCmAmGmAmUmUmA/3ZEN/ |

**Primers** were ordered as custom DNA oligos (Integrated DNA Technologies) at 25 nmol scale.

| | |
|---|---|
| CD44 3'UTR TS Forward | ACTCGAGACCAAAGGTTTTCCATCCTGTCC |
| CD44 3'UTR TS Reverse | AGCGGCCGCACATCTCTCCTTTAAAGATATTCTAC |
| CDK12 3'UTR TS Forward | ACTCGAGAcaaaaacctttcaaacagagc |
| CDK12 3'UTR TS Reverse | AGCGGCCGCActgttcttcctcacagtatgc |
| sdRNA-D19b qRT-PCR | GCCCATTACAAGATCCAACTCTGAT |
| sdRNA-A24 qRT-PCR | GCTCCATGTATCTTTGGGACCTGTCA |
| U6 Forward | GCTCGCTTCGGCAGCACATATAC |
| U6 Reverse | CGCTTCACGAATTTGCGTGTCATCC |
